# Sulfate import in *Salmonella* Typhimurium impacts bacterial aggregation and the neutrophil respiratory burst

**DOI:** 10.1101/2020.11.06.372433

**Authors:** TL Westerman, MK Sheats, JR Elfenbein

## Abstract

During enteric salmonellosis, neutrophil generated reactive oxygen species alter the gut microenvironment favoring survival of *Salmonella* Typhimurium. While the type-3 secretion system-1 (T3SS-1) and flagellar motility are potent *Salmonella* Typhimurium agonists of the neutrophil respiratory burst *in vitro*, neither of these pathways alone are responsible for stimulation of a maximal respiratory burst. In order to identify *Salmonella* Typhimurium genes that impact the magnitude of the neutrophil respiratory burst, we performed a two-step screen of defined mutant libraries in co-culture with neutrophils. We first screened *Salmonella* Typhimurium mutants lacking defined genomic regions, followed by the individual mutants mapping to genomic regions under selection. Mutants in four genes, *STM1696* (*sapF*), *STM2201* (*yeiE*), *STM2112* (*wcaD*), and *STM2441* (*cysA*), induced an attenuated respiratory burst. We linked the altered respiratory burst to reduced T3SS-1 expression and/or altered flagellar motility for two mutants (Δ*STM1696* and Δ*STM2201*). The Δ*STM2441* mutant, defective for sulfate transport, formed aggregates in minimal media and adhered to surfaces in rich media, suggesting a role for sulfur homeostasis in regulation of aggregation/adherence. We linked the aggregation/adherence phenotype of the Δ*STM2441* mutant to biofilm-associated protein A and flagellins and hypothesize that aggregation caused the observed reduction in the magnitude of the neutrophil respiratory burst. Our data demonstrate that *Salmonella* Typhimurium has numerous mechanisms to limit the magnitude of the neutrophil respiratory burst. These data further inform our understanding of how S*almonella* may alter neutrophil antimicrobial defenses.

## Introduction

Non-typhoidal *Salmonella* (NTS) infections account for the largest disease burden of foodborne illnesses globally [1]. Acute enteric disease caused by NTS is characterized by marked neutrophilic enteritis and inflammatory diarrhea [2]. Neutrophil influx into the gut lumen is induced by effector proteins secreted by the type-3 secretion system-1 (T3SS-1) encoded within the horizontally-acquired locus, *Salmonella* Pathogenicity Island-1 [3]. Neutrophilic influx benefits NTS by eliminating competing microbes and generating new nutrients to support NTS metabolism in the gut lumen [4–6]. However, neutrophils also limit the systemic spread of NTS in the host [7]. Given this, it is apparent that there is a complicated interplay between NTS and neutrophils that impact NTS fitness during infection.

A hallmark of the neutrophil antimicrobial response is the production of reactive oxygen species (ROS) through the respiratory burst [8, 9]. Generation of ROS occurs through assembly of the multicomponent NADPH oxidase enzyme complex at the neutrophil phagolysosome and/or cell membrane, with ROS released intracellularly, extracellularly, or both [10, 11]. The magnitude of the neutrophil respiratory burst is influenced by interactions with the invading pathogen. Lipopolysaccharide, a pathogen associated molecular pattern located in the bacterial outer membrane, is one factor that primes the neutrophil for a robust respiratory burst [12, 13]. In addition, both the *Salmonella* Typhimurium T3SS-1 and flagellar motility are important agonists of the neutrophil respiratory burst *in vitro* [14]. However, biofilm-associated NTS, encased in an extracellular matrix composed of proteins, carbohydrates, and extracellular DNA (reviewed in [15]), elicit a diminished neutrophil respiratory burst when compared to planktonic cells [16]. Taken together, these findings suggest that NTS have numerous mechanisms to alter the magnitude of the neutrophil respiratory burst, which likely help to facilitate its survival in different host compartments.

The mechanisms by which NTS modulate the respiratory burst are unknown. Although the T3SS-1 and flagellar motility are key *Salmonella* Typhimurium agonists of the neutrophil respiratory burst, deletion of both elements does not completely abrogate the neutrophil respiratory burst *in vitro* [14]. Therefore, we hypothesized that additional *S*. Typhimurium factors would serve as agonists of the neutrophil respiratory burst. Using a two-step genetic screen, we identified genes that were associated with significant changes in the neutrophil respiratory burst and characterized the genes for effects on T3SS-1 expression and swimming motility. We found that cellular aggregation mediated by a defect in sulfate import reduced the magnitude of the neutrophil respiratory burst. Together this work provides mechanistic insight into how *S*. Typhimurium modulates the magnitude of the neutrophil respiratory burst, a phenomenon that likely facilitates its survival during enteric infection.

## Methods

### Bacterial strains and growth conditions

All bacterial strains are derivatives of *Salmonella enterica* serovar Typhimurium ATCC 14028s. The multigene deletion (MGD) and single-gene deletion (SGD) mutant library collections are previously described [17]. Mutations were moved into a clean genetic background by P22 transduction and antibiotic cassettes were removed as described [18, 19]. Unless otherwise indicated, bacteria were grown in Luria Bertani (LB-Miller) broth or LB agar supplemented with nalidixic acid (50 mg/L), kanamycin (50 mg/L), chloramphenicol (20 mg/L), carbenicillin (100 mg/L), and tetracycline (20 mg/L). For assays using bacteria from stationary phase growth, bacteria were grown in LB broth at 37°C with agitation (225 rpm) overnight. For assays using bacteria from late-exponential growth, bacteria were prepared by diluting overnight cultures 1:100 in LB broth followed by incubation at 37°C with agitation for 3.5 hours.

For growth curves, overnight cultures were diluted 1:100 into 50 mL LB or M9 minimal media and grown at 37°C with agitation (225 rpm) for 24 hours. M9 minimal media (48 mM Na_2_HPO_4_, 22 mM KH_2_PO_4_, 9 mM NaCl, 19 mM NH_4_Cl, 0.1 mM CaCl_2_ and 2 mM MgSO_4_) was supplemented with 0.2% (w/v) dextrose as a carbon source and 1 mM L-cysteine, where indicated [20]. Samples were taken hourly for 6 hours and once at 24 hours, diluted, and plated to enumerate colony forming units (CFU). Each experiment was performed on three independent occasions.

For growth in neutrophil-*Salmonella* co-culture media, bacteria were washed and diluted in PBS, and added to 100 μL RPMI-1640 (with L-glutamine, without phenol red; Gibco) supplemented with 1 mM Ca^++^, 1 mM Mg^++^, and 10% normal human serum from male AB donors (Corning) to achieve a final concentration of 5×10^6^ CFU. Cultures were incubated standing at 37°C with 5% CO_2_ for 2 hours and then serially diluted and plated to enumerate CFU. Each experiment was performed on three independent occasions.

For analysis of the effects of sulfur on bacterial viability, a sulfur-free (SF) minimal media was used. SF media was a modification of M9 minimal media using 2 mM MgCl_2_ as a magnesium source. SF media was supplemented with 0.5 mM sodium tetrathionate (Na_2_S_4_O_6_), 1 mM sodium thiosulfate (Na_2_S_2_O_3_), or 1 mM L-cysteine as indicated. Overnight cultures were diluted 1:100 into SF media and grown at 37°C with agitation for 24 hours. Bacterial viability was assessed at 24 hours by serial dilution and plating to enumerate CFU. The number of generations was established as (log_10_(final CFU)-log_10_(start CFU))/0.301. Each experiment was performed on three independent occasions.

### Human subjects

Neutrophils were isolated from peripheral blood of healthy adult volunteers. All participants provided written informed consent. The study was approved by the Institutional Research Ethics Committee of North Carolina State University (IRB approval #616).

### Neutrophil respiratory burst in co-culture with *Salmonella*

Neutrophils (PMN) were isolated from whole blood by a Ficoll gradient centrifugation technique as previously described [14]. Isolated PMNs exhibited greater than 98% viability and 95% purity as determined by trypan blue exclusion. Purified PMNs were suspended in RPMI-1640 (with L-glutamine, without phenol red; Gibco) supplemented with 1 mM Ca^++^, 1 mM Mg^++^, and 10% normal human serum from male AB donors (Corning) to a final concentration of 1.15×10^6^ PMN/mL. Neutrophils were primed with human recombinant granulocyte-macrophage colony stimulating factor (GMCSF; 30 ng/mL) for 30 minutes at 37°C with 5% CO_2_.

For library screening, bacteria from stationary phase growth were diluted 1:1 in phosphate buffered saline (PBS) prior to addition to the neutrophil culture. For individual mutant strain evaluation, bacteria from stationary and late-exponential growth phases were used as indicated to inoculate neutrophil cultures. Bacteria were washed and diluted in PBS to achieve a multiplicity of infection (MOI) of 50:1. Bacteria were serially diluted and plated on LB agar to establish the final MOI.

Intracellular ROS was measured by fluorescence of rhodamine generated by oxidation of dihydrorhodamine-123 (DHR-123) as previously described [14]. Neutrophils were allowed to settle for 10 minutes on black polystyrene 96-well plates coated with 5% FCS prior to addition of 10 μM DHR-123 and either bacteria or controls. Negative controls in each plate included neutrophils with assay media (RPMI + 10% human serum) and bacterial growth media (LB) to ensure neutrophils were not inappropriately activated. Neutrophils stimulated with phorbol 12-myristate 13-acetate (PMA; 50 ng/mL; Sigma Aldrich) were included on each plate as a positive control. A mutant we previously determined to have an altered respiratory burst (ΔSPI-1) was also included on each plate as a control for the assay [14]. Plates were incubated at 37°C with 5% CO_2_. Relative fluorescence units (RFU; 485 nm excitation, 528 nm emission) was determined prior to incubation and then hourly for 3 hours for library screening, and every 30 minutes for 2 hours for individual strain evaluation (Synergy HTX, BioTek). Neutrophils from 2 different blood donors were used for library screens and neutrophils from 3-4 different blood donors were used for individual strain evaluation.

Total intracellular and extracellular ROS was measured by luminol (5-amino-2,3-dihydro-1,4-phthalazinedione; 1 mM; Sigma-Aldrich) enhanced chemiluminescence as previously described [14]. Briefly, neutrophils were aliquoted onto a white polystyrene 96-well plate coated with 5% FCS. Neutrophils were allowed to settle for 10 minutes prior to addition of luminol and a baseline luminescence measurement was obtained (integration time 1s; Synergy HTX, BioTek). Neutrophils were then stimulated with bacteria or PMA as a positive control. The plate was incubated at 37°C and luminescence measured every 5 minutes for 90 minutes. The peak and time to peak luminescence were determined for each strain and normalized to the WT where indicated.

### Bacterial motility assays

Swimming motility was assayed on plates containing 0.3% Difco Bacto Agar (LB Miller base, 25 g/L) as described [21]. Overnight cultures were grown at 37°C with agitation and cell concentration was normalized by OD_600_. Bacteria were spotted onto plates and incubated at 37°C for 4 hours. The diameter of each colony was measured and compared to the wild-type organism on the same plate. Each assay was performed on three separate occasions in 4-5 replicates. Swimming assays were repeated on individual plates for photographs (ChemiDoc MP).

### ß-galactosidase assays

Bacteria bearing plasmid constructs were grown overnight at 37°C with agitation in LB broth supplemented with the appropriate antibiotic. For induction of T3SS-1 expression, overnight cultures were diluted 1:100 into LB with antibiotic and incubated at 37°C with agitation for 3.5 hours. ß-galactosidase activity was determined from cell pellets using standard methodology and Miller Units determined using the following equation: 1000 x [OD_420_ – (1.75 x OD_550_)] / [time x volume x OD_600_] [22]. Experiments were performed on five separate occasions.

### Complementing plasmid construction

Genomic DNA was isolated from *S*. Typhimurium using the GenElute bacterial genomic DNA kit (Sigma-Aldrich). Restriction endonuclease sequences were incorporated into the following primer sequences to facilitate cloning: STM2441BamH1Fwd 5’ GTAGGATCCGCGATTTTACTGGGCGCATC 3’ and STM2441Kpn1Rev 5’ GTAGGTACCTGTAATTTGACCAGCGGCGT 3’. A 1.5kb product for *STM2441* including the native promoter was generated from template genomic DNA using Q5 DNA polymerase (New England Biolabs) with an annealing temperature of 72°C and an extension time of 40s for 35 cycles. The expected size of the PCR product was confirmed by agarose gel electrophoresis. The PCR product was digested with the restriction endonucleases BamH1 and Kpn1 (New England Biolabs) and purified (QIAquick® PCR Purification Kit, Qiagen). The low-copy number vector, pWSK29 [23], was sequentially digested with BamH1 and Kpn1 followed by dephosphorylation with shrimp alkaline phosphatase (New England Biolabs). Ligation of the product and linearized vector was performed overnight at 14°C with T4 DNA ligase (New England Biolabs). The resulting construct was transformed into DH5α *Escherichia coli* by heat shock and transformants were obtained by selection on LB agar with carbenicillin and X-gal (5-bromo-4-chloro-3-indolyl-β-D-galactopyranoside, 40 μg/mL). Transformants were purified twice and plasmid isolated (QIAprep® Miniprep Kit, Qiagen). The insert size length was determined by restriction digestion of the plasmid followed by agarose gel electrophoresis and the expected sequence was confirmed by Sanger sequencing (Eton Bioscience). The complementing plasmid was transformed into restriction-deficient modification-positive *S*. Typhimurium LB5000 by electroporation and transformants were isolated by selection on LB agar with carbenicillin and purified twice [24]. Plasmids were then transformed into the mutant by electroporation. The resulting strains were purified twice and then stored in glycerol stocks at −80°C.

### Crystal violet assays

Bacteria were grown to both stationary and late-exponential phase in 50 mL conical tubes in 5 mL volume. Cell density was determined by OD_600_. Non-adherent cells were removed by decanting and tubes were washed twice with ddH_2_O. Adherent cells were stained with 0.1% crystal violet for 15 minutes followed by 4 washes with ddH_2_O. Tubes were inverted and allowed to dry overnight. The crystal violet was solubilized in 30% acetic acid by vortexing and quantified by measurement of OD_550_.

### Statistical analyses

For MGD library screening, an altered respiratory burst was determined using the raw rhodamine RFU data, comparing each mutant to the WT control on the same 96-well plate by a two-way ANOVA with a Dunnet’s test for multiple comparisons. For screening the SGD mini-library, we normalized the RFU elicited by SGD mutants to the RFU elicited by the WT on the same 96-well plate at a given time to reduce bias between plates. Statistical significance was determined by two-way ANOVA with a Dunnet’s test for multiple comparisons. For all other assays, statistical significance was determined using Student’s *t*-test or two-way ANOVA with Dunnet’s correction for multiple comparisons where indicated. Significance was set at P<0.05. Analyses were performed using GraphPad Prism version 8.0.

## Results

### Library screen for *S*. Typhimurium factors that alter the neutrophil respiratory burst

Flagellar motility and the T3SS-1 are potent *S*. Typhimurium agonists of the neutrophil respiratory burst [14]. To identify additional mechanisms by which *S*. Typhimurium influences the neutrophil respiratory burst, we performed a two-step mutant library screen using defined libraries of *S*. Typhimurium mutants [17]. The multigene deletion (MGD) library consisted of 151 defined mutants, each containing deletions of ~4-40 contiguous genes, with total library coverage of nearly half of the *S*. Typhimurium genome. Individual mutants from the MGD library were co-cultured with neutrophils to establish mutants that alter the magnitude of the intracellular neutrophil respiratory burst. A mutant was considered to cause an altered neutrophil respiratory burst if it was significantly different from the WT at any time after co-culture (**Supplemental Table 1**). We identified 45 MGD mutants that elicited an altered the respiratory burst, with 2 mutants covering overlapping genomic regions (**Supplemental Table 2**). Three mutants deleted for the T3SS-1 and its effectors and 3 mutants deleted for flagellar components elicited a reduced respiratory burst in our screen, confirming our ability to identify known agonists of the neutrophil respiratory burst. Four MGD mutants covered 25 genes or more and were excluded from further analysis due to the likelihood of additive effects of multiple genes affecting the respiratory burst. We confirmed the mutant identity for 19 of 21 mutants by PCR and prioritized the genes from 19 confirmed genomic regions for further analysis. A mini-library of 134 single-gene deletion (SGD) mutants from the prioritized genomic regions was assembled.

In the second step of our screen, we interrogated the effects of our mutant mini-library on the neutrophil respiratory burst, as measured by rhodamine fluorescence. We found 42 mutants from 16 genomic regions that elicited an altered respiratory burst as compared to the WT organism (**Supplemental Tables 3 and 4**). We selected seven of the 42 SGD mutants for further evaluation in PMN-*Salmonella* co-culture. Seven mutations, in Δ*STM0018*, Δ*STM1213*, Δ*STM1690*, Δ*STM1696*, Δ*STM2112*, Δ*STM2201*, and Δ*STM2441*, were moved into a clean genetic background and re-tested in co-culture with neutrophils. Four mutants (Δ*STM1696*, Δ*STM2112*, Δ*STM2201*, and Δ*STM2441*) elicited a significantly attenuated neutrophil respiratory burst as compared to the WT organism when grown to both stationary and exponential phase growth, while three mutations had no effect on the neutrophil respiratory burst (**Figure 1**).

**Figure 1:**
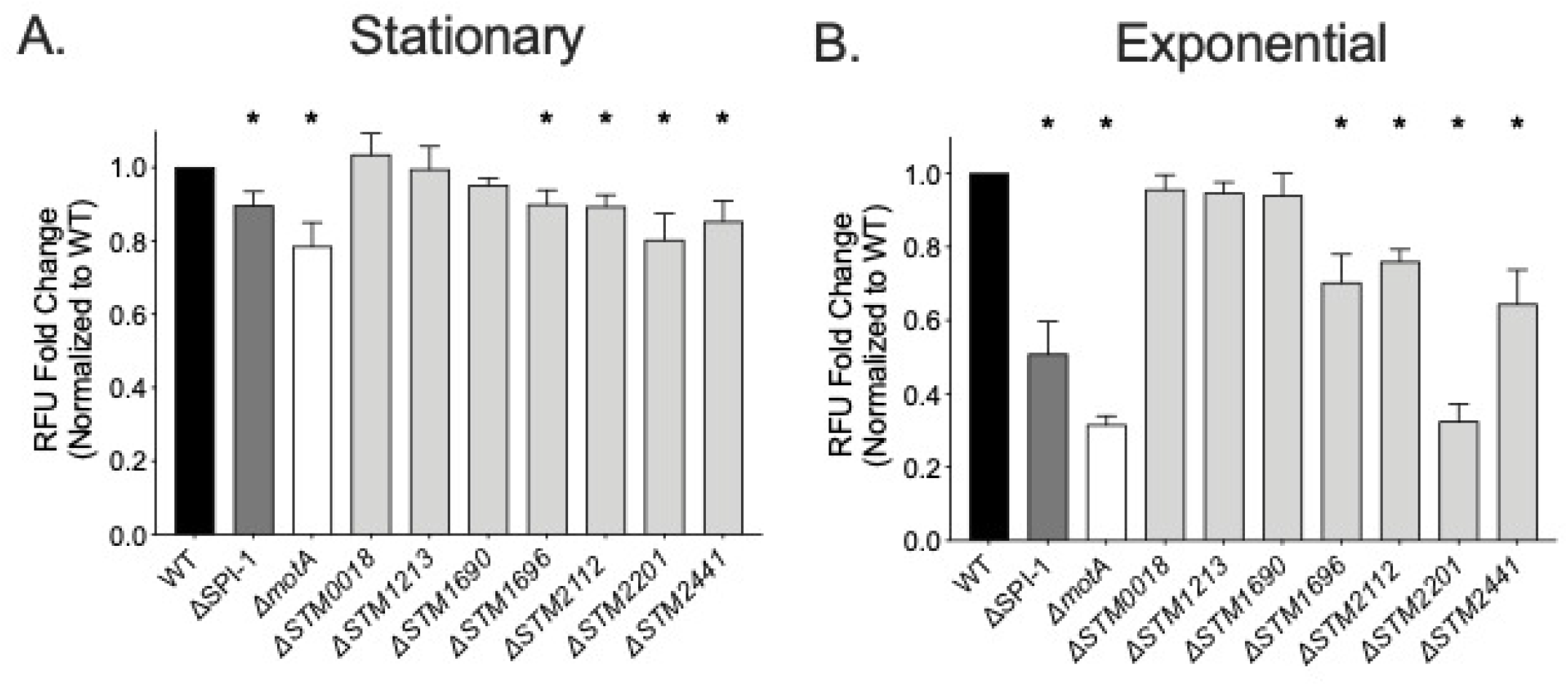
The neutrophil respiratory burst is blunted in response to four single gene deletion mutants. The neutrophil respiratory burst as measured by rhodamine fluorescence elicited by the WT (HA420) and the ΔSPI-1 (JE598), Δ*motA* (JE1202), Δ*STM0018* (JE963), Δ*STM1213* (JE965), Δ*STM1690* (JE967), Δ*STM1696* (JE969), Δ*STM2112* (JE971) Δ*STM2201* (JE973), and Δ*STM2441* (JE975) mutants at an MOI of 50:1. Neutrophils were exposed for (A) 2-hours to bacteria grown to stationary phase or (B) 1-hour to bacteria grown to late-exponential phase. The relative fluorescence elicited by each mutant was normalized to the WT organism for each donor. Bars indicate mean +/- SEM from triplicate samples using blood from 4 different donors. * indicates significant difference between the WT and indicated mutant by Student’s *t*-test with P<0.05.

The phenotypes of two of our mutants, in Δ*STM1696* and Δ*STM2201*, were reversed when tested in the clean genetic background (**Supplemental Table 4**). Overall, our screening strategy identified 4 mutants that stimulated an altered neutrophil respiratory burst as compared with the WT organism.

### Effects of mutations on swimming motility and T3SS-1 expression

We investigated possible mechanisms for the alteration in the neutrophil respiratory burst elicited by the four identified mutants. First, we performed swimming assays to determine the contribution of each gene to flagellar motility. We found a significant defect in swimming motility for the Δ*STM1696*, Δ*STM2201*, and Δ*STM2441* mutants (**Figure 2**). The Δ*STM2201* mutant was amotile. Since the magnitude of the respiratory burst stimulated by the Δ*STM2201* mutant was comparable to the amotile Δ*motA* mutant, it is likely that the severe motility defect of the Δ*STM2201* mutant explains the observed alteration in the neutrophil respiratory burst.

**Figure 2:**
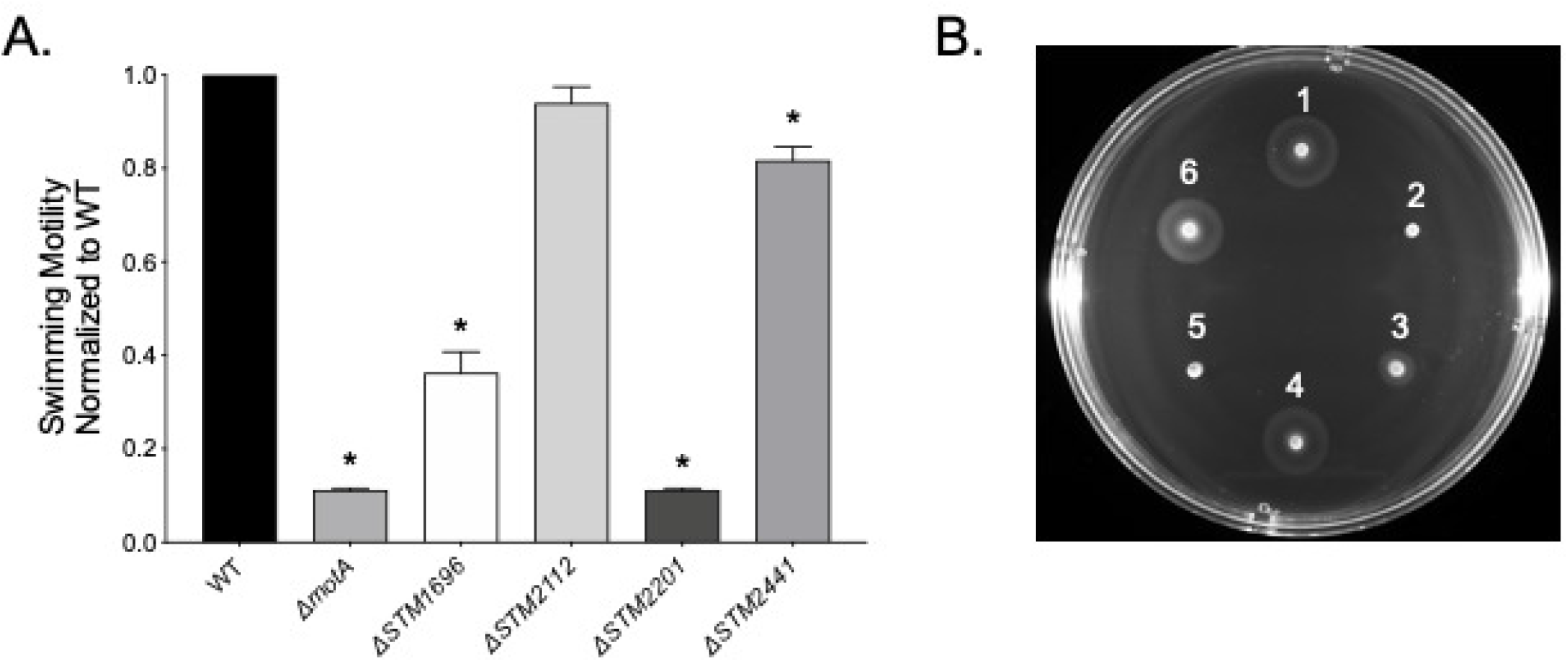
The Δ*STM1696*, Δ*STM2201*, and Δ*STM2441* mutants have reduced swimming motility. (A) Normalized overnight cultures of the WT and the Δ*motA* (JE1202), Δ*STM1696* (JE969), Δ*STM2112* (JE971) Δ*STM2201* (JE973), and Δ*STM2441* (JE975) mutants were spotted onto swimming plates. Cell spread was measured 4 hours post-inoculation. Each assay was performed in 4-5 replicates on 3 separate occasions. Bars represent mean +/- SEM. * indicates significant difference between the WT and mutant by Student’s *t*-test with P<0.05 (B) Representative photograph of a swimming plate (1: WT; 2: Δ*motA*; 3: Δ*STM1696*; 4: Δ*STM2112*; 5: Δ*STM2201*; 6: Δ*STM2441*).

Next, we evaluated the effects of our mutations on expression of the T3SS-1 using a plasmid containing a T3SS-1 apparatus gene promoter (P_*prgH*_) driving the expression of *lacZY*. We found reduced T3SS-1 expression for the Δ*STM1696* mutant only (**Figure 3**). The combination of decreased motility and altered T3SS-1 expression for the Δ*STM1696* mutant suggests that one or both of the observed defects contribute to the weakened neutrophil respiratory burst. Through these experiments evaluating both swimming motility and T3SS-1 expression, we were able to assign likely causes for the observed defects in neutrophil respiratory burst for both the Δ*STM1696* and Δ*STM2201* mutants.

**Figure 3:**
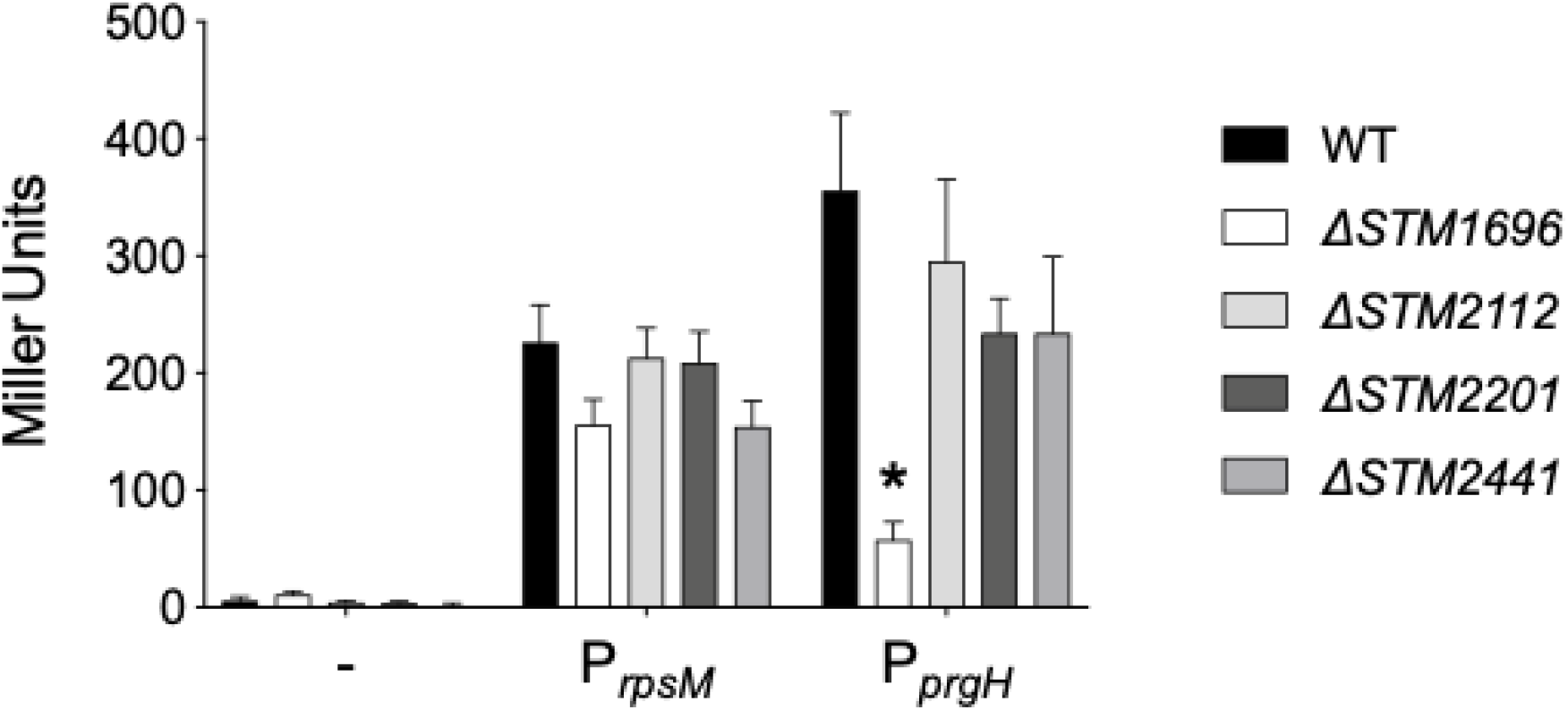
Expression of the T3SS-1 is reduced in the Δ*STM1696* mutant. Activation of a T3SS-1 apparatus promoter (P_*prgH*_) in late-exponential growth as determined by ß-galactosidase activity. Plasmid constructs with the indicated promoter driving *lacZY* expression included: – (promoter-less *lacZY*), P_*rpsM*_ (positive control), and P_*prgH*_ (T3SS-1 promoter). Bars represent the mean +/- SEM from assays performed on five separate occasions. Statistical significance (*) was determined by Student’s *t*-test with P<0.05.

### Colanic acid and the neutrophil respiratory burst

Colanic acid is a negatively charged polysaccharide capsule that helps to maintain transmembrane potential and the proton motive force during envelope stress [25]. Colanic acid is produced by enzymes encoded within the colanic acid capsule biosynthetic gene cluster (*STM2118-STM2099*) and its production is stimulated by low temperatures (<30°C) and biofilm forming conditions [26–28]. Since colanic acid is not typically produced in rich media at body temperature, we found it surprising that two mutants in the biosynthetic gene cluster, Δ*STM2112* (Δ*wcaD*) and Δ*STM2114* (Δ*wcaB*) were identified in the screen.

To establish whether colanic acid acts as an agonist of the neutrophil respiratory burst in our assay conditions, we investigated the neutrophil respiratory burst in response to a complete colanic acid capsule cluster mutant (Δ*wza*), deleted for *wza* through *wcaM* (*STM2118-STM2099*) [25]. We observed no alterations to the neutrophil respiratory burst for the Δ*wza* mutant in comparison to the isogenic WT (**Figure 4**). Since mutants within the colanic acid biosynthesis pathway located downstream of the WcaJ glycosyl transferase could accumulate toxic intermediates that affect viability, we evaluated the growth of the Δ*STM2112* mutant in different media [29]. We found no growth defects in either rich or minimal media (**Supplemental Figure 1a-b**). Neutrophil-*Salmonella* co-culture media includes 10% normal human serum and therefore contains complement. Colanic acid protects *E. coli* against complement-mediated killing [30], therefore we hypothesized the Δ*STM2112* mutant would be sensitive to the serum in our assay. We found no growth defect for the Δ*STM2112* mutant after 2 hours, the duration of time used for our neutrophil-*Salmonella* co-culture (**Supplemental Figure 1c**). Our data suggest that a defective colanic acid capsule was not the likely cause for the reduced neutrophil respiratory burst elicited by the Δ*STM2112* mutant in the conditions tested.

**Figure 4:**
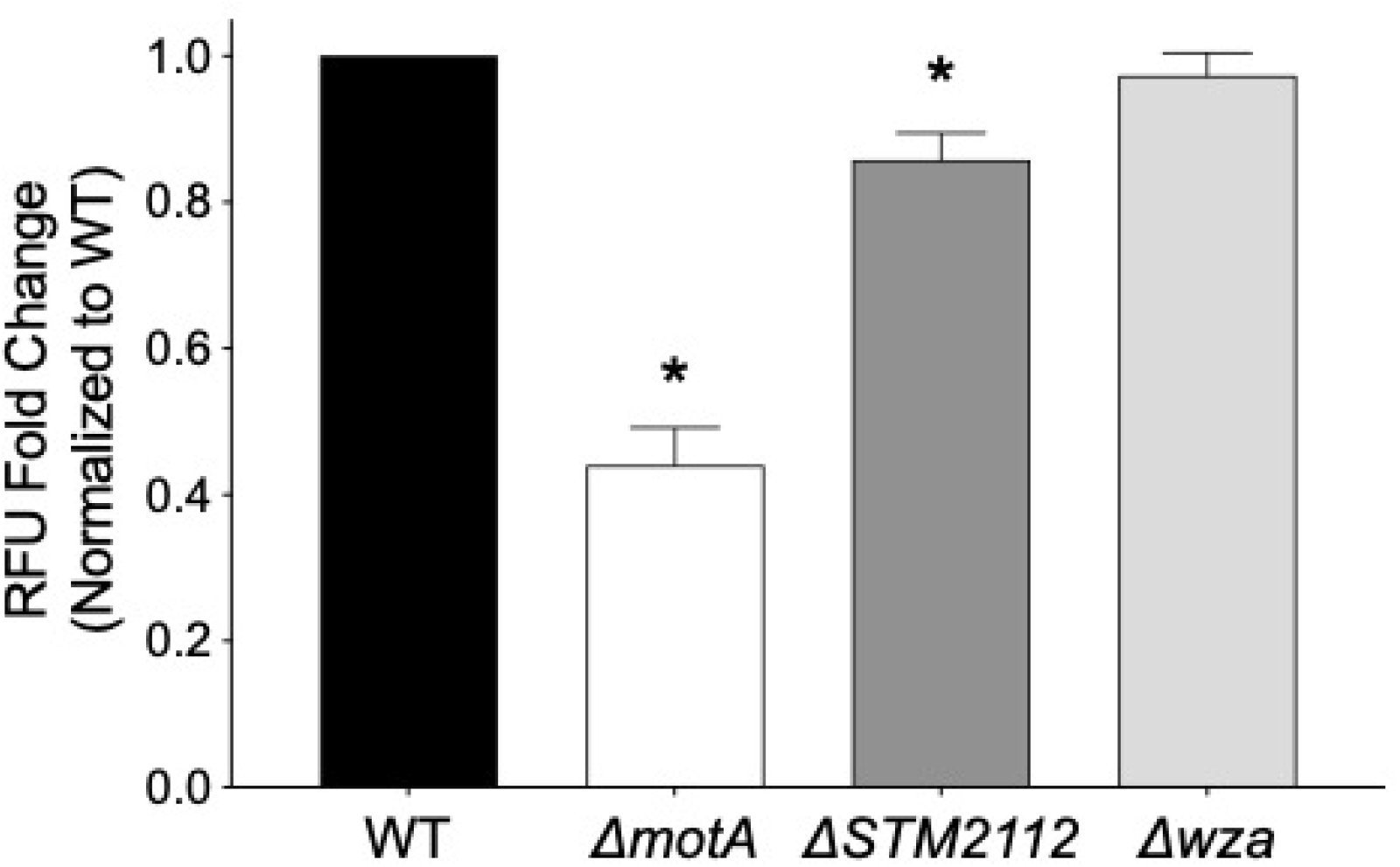
Deletion of the complete colanic acid biosynthetic cluster has no effect on the neutrophil respiratory burst. Neutrophils were exposed to the WT (WN150) or the Δ*motA* (JE1762), ΔSPI-1 (JE1760), Δ*STM2112* (JE1800), and Δ*wza* (JP245) mutants grown to late-exponential phase and the neutrophil respiratory burst measured by rhodamine fluorescence after 1-hour co-culture. Data analysis as in Figure 1. Bars indicate mean +/- SEM from triplicate samples using blood from 3 different donors. * indicates significant difference between the WT and mutant by Student’s *t*-test with P<0.05.

### Sulfate import stimulates the neutrophil respiratory burst

*STM2441* (*cysA*) encodes for the ATPase subunit of the SulT sulfate permease responsible for importing sulfate and thiosulfate into the cell [31]. Our data demonstrate that the Δ*STM2441* mutant elicited a lower magnitude neutrophil respiratory burst as compared to the WT (**Figure 1**). We constructed a low-copy number plasmid containing an intact copy of *STM2441* with its native promoter to evaluate the phenotypes of the mutant complemented *in trans*.

Complementation of the Δ*STM2441* mutant *in trans* completely restored the intracellular neutrophil respiratory burst to WT levels, definitively linking the altered respiratory burst to deletion of *STM2441* (**Figure 5a**). To verify our observations, we tested the effect of the Δ*STM2441* mutant on the neutrophil respiratory burst using luminol, which measures both intracellular and extracellular reactive oxygen species [32]. We found that the peak magnitude of the neutrophil respiratory burst elicited by the Δ*STM2441* mutant was decreased when compared with the WT organism (**Figure 5b** and **Supplemental Figure 2**), but there was no effect of the mutation on the time to peak respiratory burst (**Figure 5c** and **Supplemental Figure 2**). Finally, we found no reversal of the swimming defect for the complemented Δ*STM2441* mutant (**Figure 5d-e**). Together our data demonstrate that neutrophils produce a muted respiratory burst in response to the Δ*STM2441* mutant, a phenotype that is not likely related to the small reduction in swimming motility.

**Figure 5:**
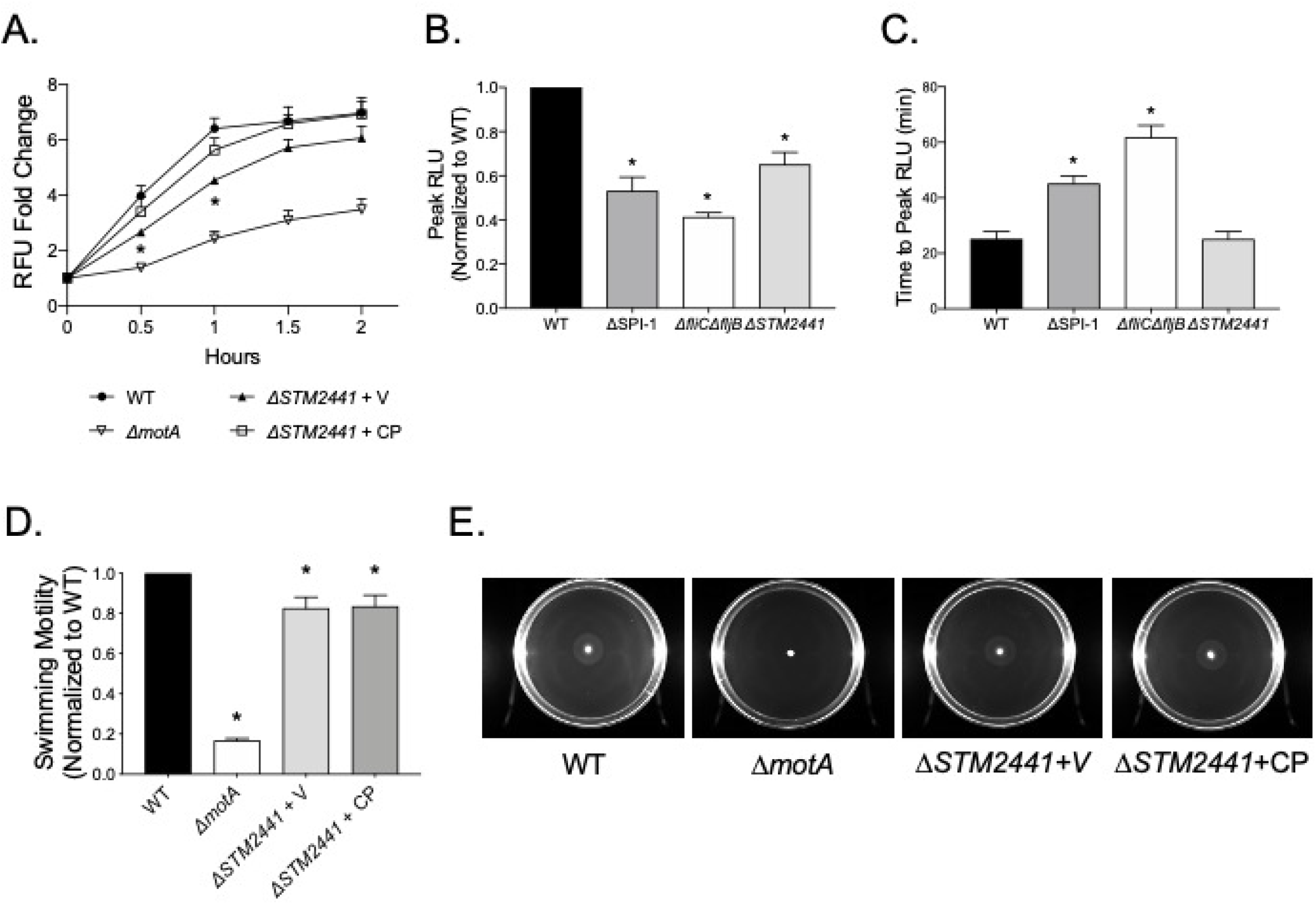
Deletion of Δ*STM2441* is linked to altered total and intracellular neutrophil respiratory burst. Neutrophils were exposed to the WT or the Δ*motA* (JE1202), Δ*STM2441* (JE975), Δ*fliC*Δ*fljB* (JE524), Δ*STM2441* + V (pWSK29; JE1673), and Δ*STM2441* + CP (pWSK29::*STM2441*; JE1675) mutants grown to late-exponential phase. (A) The neutrophil respiratory burst was measured by rhodamine fluorescence as in Figure 1 or (B and C) luminol-enhanced chemiluminescence. (B) Peak luminescence normalized to the WT for the same donor and (C) time to peak luminescence were determined from each donor. Data points indicate mean +/- SEM from triplicate samples using blood from 3 different donors. (D) Normalized overnight cultures of the WT and the Δ*motA*, Δ*STM2441* + V, and Δ*STM2441* + CP mutants were spotted onto swimming plates. Analysis as in Figure 3. (E) Representative photographs of swimming plates. * indicates statistical significance between the indicated mutant and the WT by Student’s *t*-test with P<0.05.

### Sulfate import defect induces cellular aggregation

The Δ*cysA* mutant is a cysteine auxotroph because sulfate import is essential for cysteine biosynthesis [33, 34]. We investigated the growth of the Δ*STM2441* mutant in both rich and minimal media. As anticipated, we found that the Δ*STM2441* mutant has a growth defect, reversed by complementation *in trans*, in M9 minimal media which contains MgSO_4_ as the sole sulfur source (**Figure 6a**). The addition of L-cysteine to M9 minimal media completely reversed the growth defect, confirming that the Δ*STM2441* mutant is a cysteine auxotroph (**Figure 6b**). We found no growth abnormalities in rich media (LB) or media used for *Salmonella*-neutrophil co-culture, which contains 0.2 mM L-cystine and 0.4 mM MgSO_4_ (**Supplemental Figure 3a-b**).

**Figure 6:**
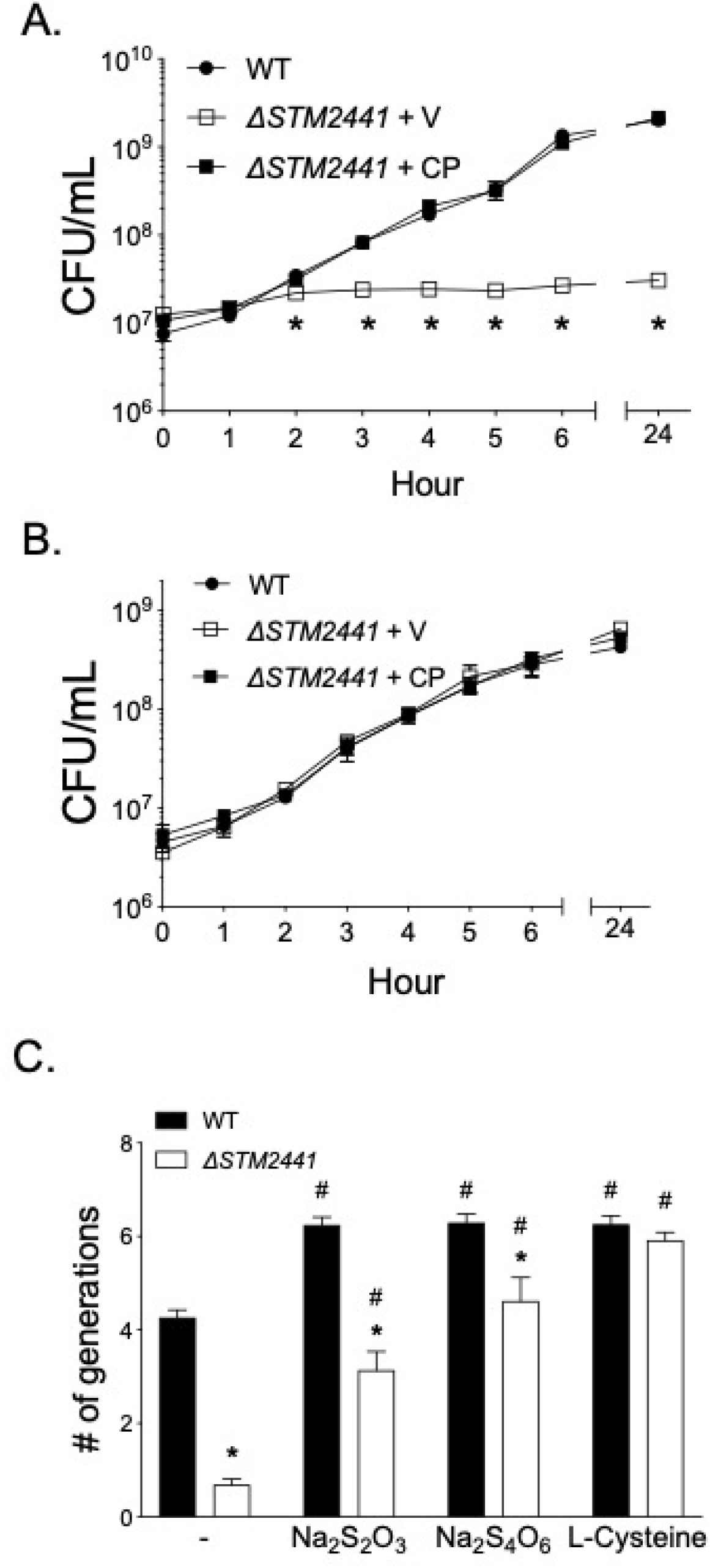
The growth defect of the Δ*STM2441* mutant cysteine auxotroph is partially reversed by thiosulfate and tetrathionate. Growth of the WT, Δ*STM2441* + V (pWSK29; JE1673), and Δ*STM2441* + CP (pWSK29::*STM2441*; JE1675) in (A) M9 minimal broth or (B) M9 with L-cysteine (1 mM). (A-B) Overnight cultures were diluted 1:100 into indicated media and samples taken hourly to determine CFU/mL. Data points represent mean +/- SEM. CFU data were log transformed and statistical significance determined by two-way ANOVA. * indicates significant difference between WT and Δ*STM2441* + pWSK29 with P<0.05. (C) Growth of the WT and Δ*STM2441* (JE975) mutant in SF broth with no sulfur source (-) or supplemented with sodium thiosulfate (1 mM Na_2_S_2_O_3_), sodium tetrathionate (0.5 mM Na_2_S_4_O_6_), or L-cysteine (1 mM). Overnight cultures were diluted 1:100 into indicated media and the number of generations was determined after 24h growth. Statistical significance was determined by Student’s *t-*test with P<0.05. * indicates significant difference between WT and Δ*STM2441* in a given media. # indicates significant difference between the indicated media and non-supplemented media for a given strain. Experiments were performed on three independent occasions.

Since sulfate and thiosulfate can be used in sulfate assimilation pathways, we hypothesized that other sulfur sources known to be present in the host could rescue the growth defect of the Δ*STM2441* mutant [4, 31, 35]. We evaluated growth in media with thiosulfate or tetrathionate as sole sulfur sources. We found that both thiosulfate and tetrathionate partially restored growth of the Δ*STM24441* mutant (**Figure 6c**). The partial restoration in growth suggests that *Salmonella* has one or more other import mechanism(s) that allow selective import of thiosulfate and/or tetrathionate. While performing experiments in minimal media, we identified macroscopic cellular aggregates within the liquid media that also adhered to the flask walls after 24 hours of growth (**Supplemental Figure 4a**). The adherent aggregates were positive for crystal violet staining and were most abundant in media supplemented with tetrathionate and thiosulfate (**Supplemental Figure 4b**). These observations led us to investigate whether cellular aggregates could form in the bacterial preparation conditions used for the neutrophil respiratory burst assay.

Bacteria prepared for neutrophil respiratory burst assays were grown in rich media to either stationary or exponential growth. Using crystal violet staining, we detected bacterial adherence to the surface of conical tubes at the liquid-air interface in bacteria grown to both stationary and exponential phases for the Δ*STM2441* mutant (**Figure 7a**), with no apparent adherence of the WT organism, as expected. The surface adhesion was reversed in the complemented Δ*STM2441* mutant (**Figure 7a**). These data suggest that defective import of sulfate induces inappropriate bacterial aggregation, a phenotype that may explain the altered neutrophil respiratory burst stimulated by the Δ*STM2441* mutant.

**Figure 7:**
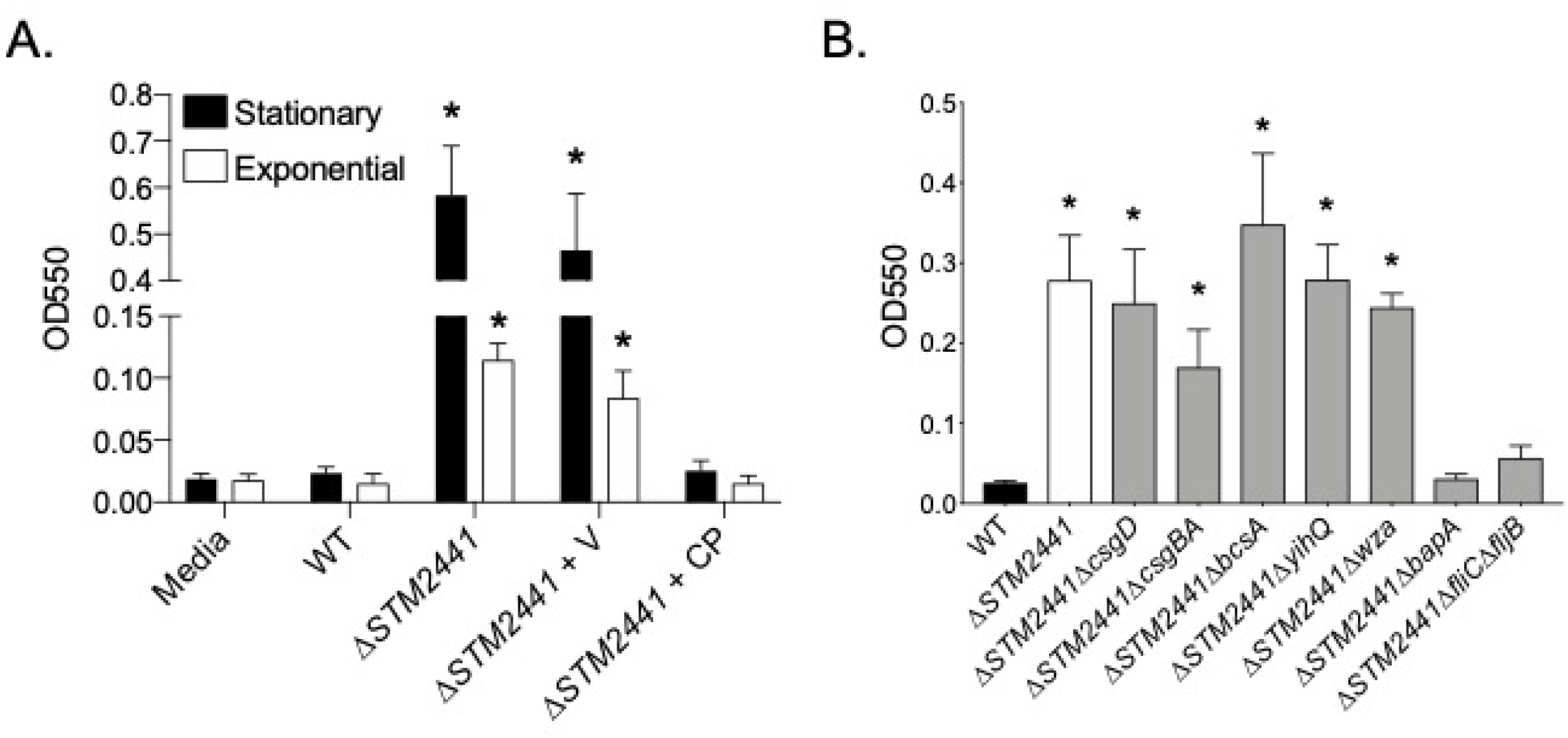
Adherence of Δ*STM2441* mutant to surfaces is linked to biofilm-associated protein and flagellins. (A) Crystal violet staining was used to quantify surface-adherent bacteria from the WT and the Δ*STM2441* mutant grown at 37°C with agitation to stationary and exponential phases. (B) Surface-adherent bacteria from the WT, Δ*STM2441* mutant (JE1391) and double mutants grown to stationary phase as determined by crystal violet staining. Crystal violet was quantified by optical density (550 nm). Experiments were performed on 4 (A) or 6 (B) independent occasions. Bars represent mean +/- SEM. (A) * indicates significant difference between the WT and mutant as determined by Student’s *t*-test with P<0.05.

We hypothesized that cellular aggregation in the Δ*STM2441* mutant was due to induction of extracellular matrix components of biofilms. Using a genetic approach, we tested whether the inappropriate aggregation observed in the Δ*STM2441* mutant aggregation could be relieved by deletion of genes involved in biofilm including *csgD* (biofilm master regulator), *csgBA* (curli fimbriae), *bcsA* (cellulose), *bapA* (biofilm associated protein A), *yihQ* (O-antigen capsule), and *wza-wcaM* (colanic acid) (reviewed in [36]). We also tested a *fliC* and *fljB* (flagellins) double mutant since flagella can be incorporated into biofilm extracellular matrix and are important for initial surface attachment [37]. The deletion of either Δ*bapA* or Δ*fliC*Δ*fljB* in the Δ*STM2441* mutant background (Δ*STM2441*Δ*bapA* and Δ*STM2441*Δ*fliC*Δ*fljB* double mutants) eliminated the adherence of the Δ*STM2441* mutant (**Figure 7b**). None of the single biofilm extracellular matrix component mutants adhered to plastic in the assay conditions used (**Supplemental Figure 5a**). Together, these data indicate that the aggregation stimulated by defective sulfate import in *S*. Typhimurium is due to both the biofilm associated protein A (BapA) and flagellins (FliC and/or FljB).

## Discussion

In this study, we used a two-step screening approach to discover new *Salmonella* Typhimurium agonists of the neutrophil respiratory burst. We first tested the effects of deletion of large genomic regions (multigene deletion mutants; MGD) on the magnitude of the neutrophil respiratory burst and followed with testing mutants in individual genes (single-gene deletion mutants) corresponding to the disrupted genomic regions identified in the first step. We identified four mutants, in Δ*STM1696*, Δ*STM2112*, Δ*STM2201*, and Δ*STM2441*, that elicited a consistently diminished neutrophil respiratory burst. We were unable to determine a mechanism for the reduction in neutrophil respiratory burst for one mutant (Δ*STM2112*). In the other three mutants, the attenuated respiratory burst response of neutrophils was linked to abnormal swimming motility (Δ*STM1696* and Δ*STM2201*), reduced T3SS-1 expression (Δ*STM1696*), and cellular aggregation (Δ*STM2441*). The cellular aggregation observed in the Δ*STM2441* mutant was linked to biofilm associated protein A and flagellins.

Our two-step genetic screen allowed us to interrogate nearly half of the S. Typhimurium genome (~1959 genes) using only 151 mutants. This screening approach has been used to identify mechanisms of cytosolic replication in epithelial cells [38], poultry colonization [39], and resistance mechanisms to an antibiofilm drug [40]. Our prior work demonstrated that expression of the T3SS-1 and flagellar motility are agonists of the neutrophil respiratory burst [14]. Mutants with deleted genes encoding for flagella and T3SS-1 apparatus and effector proteins contained within the MGD library elicited a lower neutrophil respiratory burst in the MGD screen.

Identification of known neutrophil respiratory burst agonists in by screening approach demonstrated that the methodology used could identify new genomic regions that influence the neutrophil respiratory burst. One shortcoming of our approach is the possibility that additive effects of multiple gene deletions generated false positives. Three of the 19 genomic regions that were prioritized for study in the second step did not contain single genes that altered the neutrophil respiratory burst and are considered false positive results. Another potential shortcoming of the library screening strategy was that bacteria were grown to stationary phase prior to exposure to neutrophils. The expression of many genes is altered during stationary phase as compared with exponential growth, therefore some mutants may not have been identified because the neutrophil response might only be altered when expression of a given gene is induced [41, 42]. Another shortcoming was the small sample size of two blood donors used for the library screens. Despite these shortcomings, the identification of the internal controls in this assay indicated the methodology could identify new agonists of the neutrophil respiratory burst.

*STM1696* (*sapF*) is part of the *sapABCDF* operon, which confers antimicrobial peptide (AMP) resistance in *Salmonella* by exporting AMPs extracellularly [45, 46]. We hypothesized loss of this AMP resistance may have been associated with the altered neutrophil respiratory burst by reducing viability of the Δ*STM1696* mutant after phagocytosis. We found that the Δ*STM1696* mutant has decreased flagellar motility and T3SS-1 expression which are known agonists of the respiratory burst. Although we did not rule out that defective export of AMPs by the Δ*STM1696* mutant could have contributed to the altered neutrophil respiratory burst, our data suggests that the altered expression of two important agonists of the neutrophil respiratory burst are likely cause(s) of the observed phenotype. Which of these processes is more important for the neutrophil response and why they are altered in the Δ*STM1696* mutant require further study.

The putative LysR type transcriptional regulator, *STM2201* (*yeiE*), is uncharacterized in *Salmonella* Typhimurium [43]. We correlated the diminished neutrophil respiratory burst for the Δ*STM2201* mutant to a lack of swimming motility, as neutrophils responded to the Δ*STM2201* mutant from a clean genetic background in a similar way as to the amotile Δ*motA* mutant. This phenotype is in contrast to the phenotype observed for the Δ*STM2201* mutant from the minilibrary, which elicited a greater magnitude respiratory burst than the WT. The contrasting respiratory burst responses elicited by the mutant during our screen versus in a clean genetic background demonstrates the need for rigorous confirmation of mutant phenotypes, as there can be potential confounding factors associated with mutants arrayed in a library. Although the YeiE regulon has been described in *E. coli*, it does not include any genes responsible for motility [44]. To our knowledge, this is the first report of *STM2201* (*yeiE*) playing a role in swimming motility. Further work is needed to characterize the mechanism by which *STM2201* contributes to altered swimming motility.

*STM2112* (*wcaD*) encodes a colanic acid polymerase within the colanic acid biosynthetic cluster. As an exopolysaccharide, colanic acid could act as an agonist of the neutrophil respiratory burst through stimulation of neutrophil pattern recognition receptors. However, our method of bacterial preparation was not performed in conditions known to induce colanic acid expression [26–28]. Furthermore, the lack of an altered neutrophil respiratory burst in response to the colanic acid operon deletion mutant (Δ*wza*) suggests that colanic acid deletion is not the likely cause for the altered neutrophil respiratory burst in response to the Δ*STM2112* mutant in our work. Consistent with our findings, a mutant in colanic acid biosynthesis (Δ*wcaM*) elicited no change in neutrophil respiratory burst when in planktonic growth [16]. Since mutations in enzymes downstream of the initial glycosyl transferase in the colanic acid biosynthesis pathway (WcaJ) lead to accumulation of toxic intermediates [29], we hypothesize that the Δ*STM2112* mutant has reduced viability in the presence of neutrophils, leading to the observed reduction in neutrophil respiratory burst.

We found a blunted neutrophil respiratory burst in response to the Δ*STM2441* mutant and correlated this finding to the development of cellular aggregates. *STM2441 (cysA)* encodes the ATPase subunit of the sulfate permease (*cysPTWA* and *sbp*), to import sulfate and thiosulfate into the cell for cysteine and methionine biosynthesis [31, 47]. We confirmed that the Δ*STM2441* mutant is a cysteine auxotroph and found that it forms aggregates both in sulfur-limited conditions and in rich media. Disruption of sulfate reduction, cysteine biosynthesis, and cysteine catabolism is associated with biofilm development in *E. coli* in the absence of canonical environmental cues [48–50]. Biofilm-associated *Salmonella* Typhimurium elicit a blunted neutrophil respiratory burst as compared with planktonic cells and the blunted respiratory burst is linked to curli fimbriae and O-antigen capsule [16]. Curli fimbriae are a major component of the *Salmonella* biofilm extracellular matrix and are recognized by innate immune receptors toll-like receptor-2 (TLR-2) and TLR9 and the nod-like receptor protein-3 inflammasome [51–54]. In spite of the importance of curli fimbriae in biofilm extracellular matrix, we observed no change to cellular aggregation for the Δ*STM2441* mutant when also deleted for Δ*csgBA* (curli fimbriae) or Δ*csgD* (regulator of biofilm formation). We hypothesize that the observed aggregates formed by the Δ*STM2441* mutant may have biofilm-like properties but do not represent mature biofilms; therefore, the mechanism by which they reduce the neutrophil respiratory burst may differ.

We linked both BapA and flagellins to the inappropriate aggregation of the Δ*STM2441* mutant. BapA is a large 386 kDa surface protein that is involved in initial surface attachment and promotes cell to cell interactions leading to biofilm formation at the air-liquid interface [55]. Flagellar motility also contributes to the initial bacterial attachment to surfaces during biofilm formation [56–58]. One possibility for the reduced neutrophil respiratory burst is that production of the large surface associated protein BapA could shield other *Salmonella* agonists from recognition by neutrophils. Another possibility is that BapA, which is comprised of a 27-tandem repeat of bacterial immunoglobulin-like domains, may alter the immune response [59]. For example, *S. aureus* possesses two surface-associated proteins with immunoglobulin binding repeat domains, Sbi and SpA, that block neutrophil Fc receptors in order to evade the immune response [60]. Finally, the formation of cellular aggregates by the Δ*STM2441* mutant may also effectively serve to reduce the number of individual bacteria detected by neutrophils, thereby reducing the magnitude of the respiratory burst [14]. Further investigation is needed to elucidate the mechanism(s) by which defects in sulfate acquisition lead to the formation of bacterial aggregates in *S*. Typhimurium, and how this phenotype impacts neutrophil antimicrobial responses.

Using a two-step screening approach to identify *Salmonella* mutants that influence the neutrophil respiratory burst, we confirmed four genes as important for stimulation of a maximal neutrophil respiratory burst. Two mutants had intriguing phenotypes. We found that *STM2201*, a putative Lys-R transcriptional regulator, seems to act as a regulator of flagellar motility. We also found that disruption of sulfate acquisition leads to inappropriate cellular aggregation and a resultant reduction in neutrophil respiratory burst. Further investigation into both of these genes is warranted to gain new insight into the myriad of mechanisms by which *S*. Typhimurium can influence the magnitude of the host immune response.

**Table 1:**
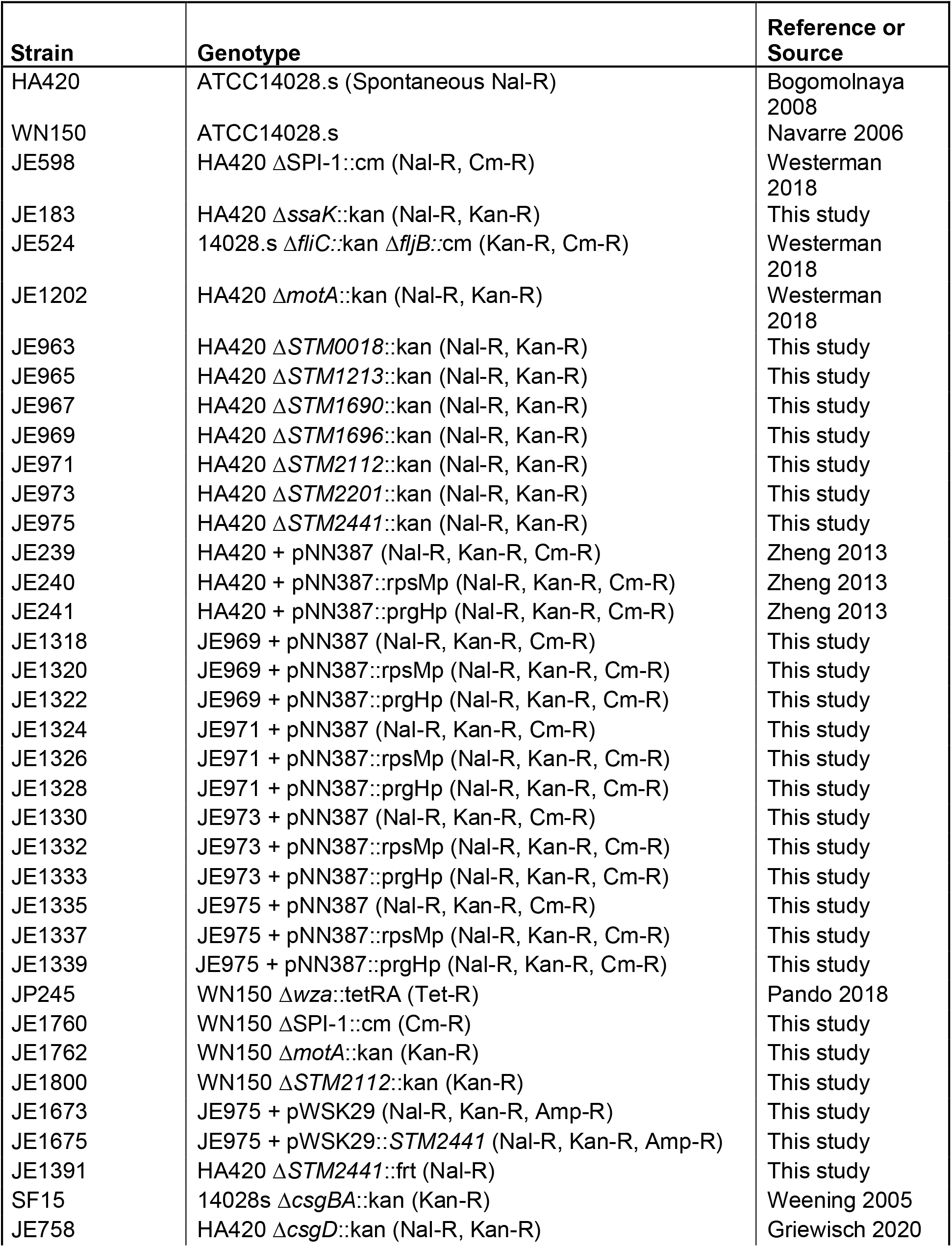
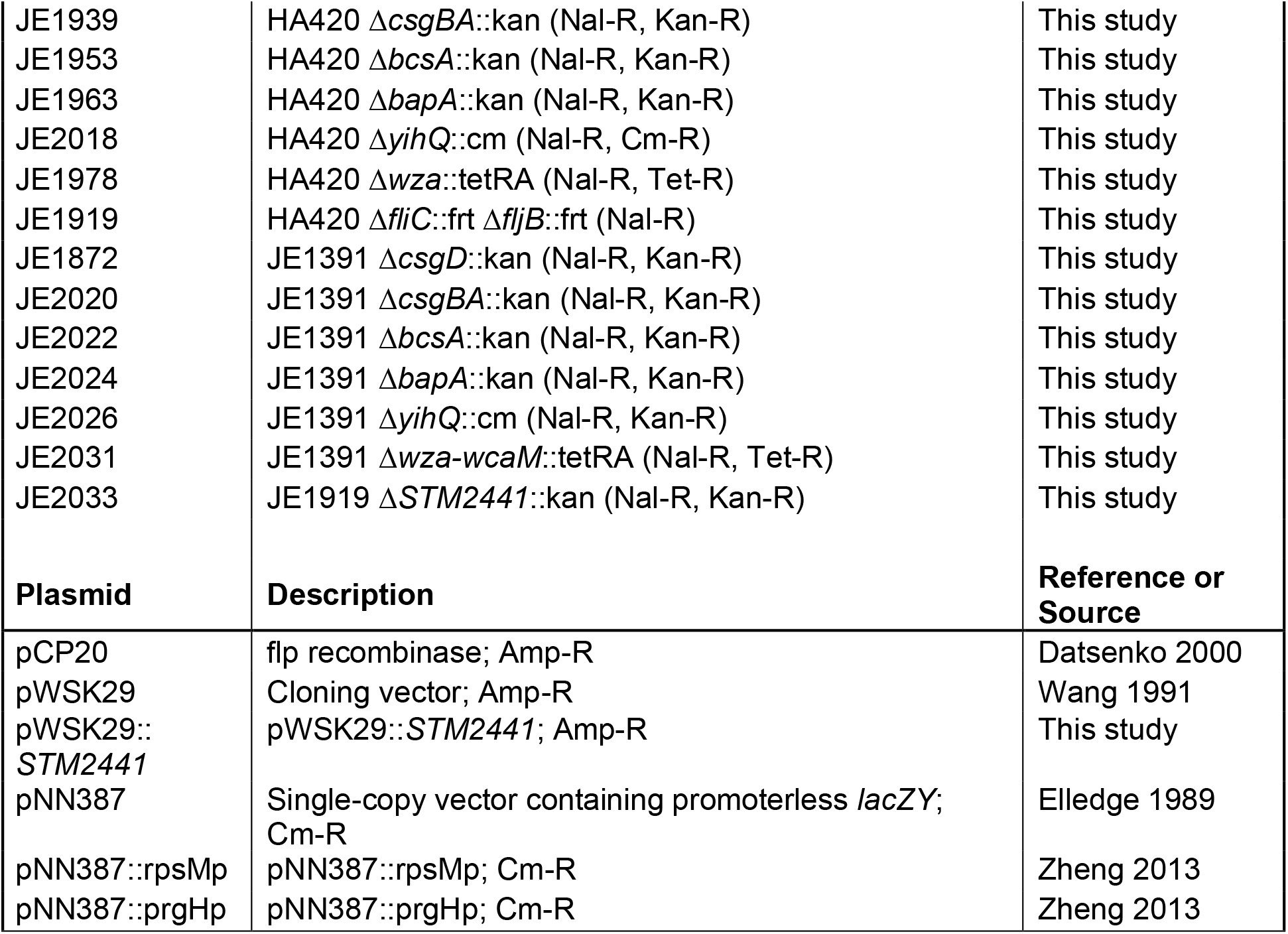
Bacterial strains and plasmids.

## Acknowledgements

A portion of this work was supported by start-up funds from the University of Wisconsin-Madison (to JRE). TLW was supported in part by 5T32OD01113. JRE was supported in part by USDA-NIFA 2018-67017-27632 and NIAID K08AI108794. MKS was supported in part by NIH Office of the Director K01OD015136. The funders played no role in the experimental design or execution of the work.

---

**Figure S1: The Δ*STM2112* mutant has normal growth in rich, minimal, and neutrophil*Salmonella* co-culture assay media.** Growth curves of the WT (HA420) and Δ*STM2112* (JE971) mutant in (A) LB or (B) M9 minimal media. Overnight cultures were diluted 1:100 into indicated media. CFU/mL were determined hourly for 6 hours and at 24 hours on three independent occasions. Data points represent mean +/- SEM. (C) Bacteria from stationary and late-exponential growth were diluted in PBS to ~5×10^6^ CFU and were added to 100 μL RPMI-1640 with 10% normal human serum in 96-well plates. Cultures were incubated standing for 2 hours at 37°C with 5% CO_2_. CFU/mL was assessed at 0 and 2 hours to determine fold growth. Bars indicate mean +/- SEM performed on three independent occasions.

**Figure S2: Total neutrophil respiratory burst as assessed by luminol-enhanced chemiluminescence from individual donors.** Bacteria from late-exponential growth were exposed to neutrophils from 3 donors. Data from individual donors was used to calculate peak luminescence and time to peak luminescence for Figure 5.

**Figure S3: The Δ*STM2441* mutant has normal growth in rich and *neutrophil-Salmonella* co-culture media.** (A) Growth curve of the WT (HA420) and Δ*STM2441* (JE975) in LB broth. (B) Growth of WT and the Δ*STM2441* mutant in neutrophil assay media as in Figure S1. Bars indicate mean +/- SEM performed on three independent occasions.

**Figure S4: Sulfur starvation induces aggregation and surface adherence of the Δ*STM2441* mutant.** Bacterial aggregates (A) and surface-adherent aggregates stained with crystal violet (B) from the Δ*STM2441* mutant grown in the indicated media. Images from bacteria prepared as for Figure 6C.

**Figure S5: Biofilm component single mutants do not adhere to surfaces.** The indicated mutants were grown as in Figure 7B and surface-adherent bacteria estimated by crystal-violet staining.

